# MULTIMODAL GRADIENTS UNIFY LOCAL AND GLOBAL CORTICAL ORGANIZATION

**DOI:** 10.1101/2024.03.11.583969

**Authors:** Yezhou Wang, Nicole Eichert, Casey Paquola, Raul Rodriguez-Cruces, Jordan DeKraker, Jessica Royer, Donna Gift Cabalo, Hans Auer, Alexander Ngo, Ilana Leppert, Christine L. Tardif, David A. Rudko, Katrin Amunts, Jonathan Smallwood, Alan C. Evans, Boris C. Bernhardt

## Abstract

Specialization of brain areas and subregions, as well as their integration into large-scale networks are key principles in neuroscience. Consolidating both local and global cortical organization, however, remains challenging. Our study developed a new approach to map global cortex-wise similarities of microstructure, structural connectivity, and functional interactions, and integrate these patterns with maps of cortical arealization. Our analysis combined repeated high-field *in-vivo* 7 tesla (7T) Magnetic Resonance Imaging (MRI) data collected in 10 healthy adults with a recently introduced probabilistic *post-mortem* atlas of cortical cytoarchitecture. We obtained multimodal eigenvectors describing cortex-wide gradients at the level of microstructural covariance, structural connectivity, and intrinsic functional interactions, and then assessed inter- and intra-area differences in cortex-wide embedding based on these multimodal eigenvectors. Inter-area similarities followed a canonical sensory-fugal gradient, with primary sensorimotor cortex being the most distinctive from all other areas, while paralimbic regions were least distinctive. This pattern largely corresponded to functional connectivity variations across different tasks collected in the same participants, suggesting that the degree of global cortical integration mirrors the functional diversity of brain areas across contexts. When studying heterogeneity within areas, we did not observe a similar relationship, despite overall higher heterogeneity in association cortices relative to paralimbic and idiotypic cortices. The results were replicated in a different dataset. Our findings highlight a close coupling between cortical arealization and global cortical motifs in shaping specialized *versus* integrative human brain function.

**Significance:** Our work situates cytoarchitecture-derived cortical areas within multimodal gradients of cortical microstructure, connectivity, and function derived from high-definition multimodal neuroimaging. We demonstrated that primary sensory and motor areas show most distinctive gradient profiles while paralimbic areas were least distinctive, overall recapitulating a sensory-fugal axis. This axis was shown to relate to the diversity of cortical areas across different functional contexts, and findings could be replicated across an independent dataset. Overall, our work shows how frameworks of cortical arealization and macroscale gradients converge in shaping functional specialization *versus* integration in the human brain.

## Introduction

Understanding how the spatial organization of the human brain gives rise to cognitive functions is a challenging, yet fundamental goal for neuroscience (1). Complex brain networks at multiple scales arise from overlapping variations in cortical microstructure, function, and connectivity (2, 3). This network involves both global integration and local specialization of cortical regions, giving rise to distributed functional communities that enable complex computations (4-7). Global integration prominently manifests within higher-order systems, notably the transmodal association cortex, which engages in increasingly abstract and self-generated cognition (8-11). In contrast, local functional specialization is more frequent in sensory and motor regions that interact more closely with the here and now (6, 8, 12, 13). The interplay between contrasting local and global motifs contributes to the hierarchical organization of the brain, underpinning both modular and integrative information processing across a diverse array of cognitive functions.

Mapping structural and functional descriptors to define discrete brain areas is essential for understanding hierarchical brain organization at macroscale. Constructing precise maps of cortical areas has been a long-standing objective in neuroanatomy, as it reduces complexity and bias when studying brain regions and inter-regional relationships (3, 14-16). Cytoarchitecture, encompassing the arrangement, distribution, composition, and layering of cells, has emerged as a gold standard to defined areas (17). Microstructural insight from this *post-mortem* approach enhances our understanding of connectivity patterns and can illuminate the role of a region in cortical functioning. However, the microstructural patterns in the human brain, and their relation to cortical function, remain challenging to address in a systematic manner due to the constraints of invasive techniques. Recently, the Julich-Brain atlas, a 3D probabilistic atlas of human brain cytoarchitecture (16) has been made available. This resource offers valuable opportunities for examining both the micro- and macro-organization of the human brain, and for contrasting local specialization with global integration. It can, thus, help to guide investigations of structure-function association across the cortical tapestry within reliably defined cortical subunits.

Gradual changes in cortical architecture have been described as well, even in early work (18). While atlas of cortical areas discretize the brain into non-overlapping constituents, recent advances emphasize a more complex organisation of the cerebral cortex, which goes beyond as simple subdivision into cortical areas, and emphasizes the role of cytoarchitectonic changes within an area (*e*.*g*., ocular dominance columns, border tuft and fringe area in the visual cortex), and changes occurring at larger scale, for example between areas (17). Moreover, recent neuroimaging has also highlighted utility of complementary descriptions of macroscale cortical organization based on eigenvector decomposition of cortex-wide similarities (commonly known as *cortical gradients* (19-21)). These gradients differentiate cortical systems in an ordered and continuous manner, and can be applied to different types of neural data. Notably, converging hierarchical trends, spanning from sensory to transmodal regions was observed across microstructural (7, 20, 22-24) and functional gradients (19, 25). These multiple dimensions can effectively capture nuanced patterns of cortical organization (23), and may provide synergy in understanding subregional heterogeneity and functional multiplicity of different cortical areas (26). More broadly, gradient mapping techniques have robustly differentiated transmodal from primary sensory/motor regions, mirroring their hierarchical contributions to cognition. In effect, such gradients have been found to align with functionally relevant properties including disproportionate expansion during primate evolution (27-29), reduced heritability and increased experience dependent plasticity (30, 31), increased network idiosyncrasy (32), and the balance of internal-vs-externally oriented processing (33-35).

Local *versus* global organization can be interrogated at the level of microstructure (e.g., cytoarchitecture), connectivity and function. In this context, MRI serves as an ideal technique to bridge structure and function across varying spatial scales (36, 37). Several contrasts have been proposed to index cortical microstructure and myelination, including T1 relaxometry (38, 39). These techniques have shown differences in intensity between primary sensory systems with marked myelination and clearly visible laminae, and heteromodal as well as paralimbic cortices with reduced myelination and less pronounced laminar differentiation (20, 40, 41). When assessing structural connectivity using diffusion MRI tractographic techniques, several studies also pointed to marked variations in connectivity profiles across cortical areas (42, 43). Notably, reports have suggested differences in the organization of region-specific connections between anterior and posterior brain regions, and between sensory/motor systems and the rest of the brain (7, 44, 45). Finally, the advent of resting-state functional MRI (rs-fMRI) has led to a surge in studies interrogating the connectivity profiles of specific regions (46-48), and delineating macroscale brain networks (4, 49-52).

Collectively, these findings underscore the versatility of MRI and its potential to conduct structure-function studies in the human brain across multiple scales. Notably, while conventional MRI acquisitions at field strengths of 3 tesla (3T) and below may have limitations in terms of resolution and signal, recent *in-vivo* studies that have moved to high fields of 7T and above have benefitted from enhanced resolution, sensitivity, and biological specificity (53-58). In addition, imaging paradigms that combine multiple MRI datasets acquired across different scanning sessions in a given individual have been shown to further increase precision for the analysis of microstructure (59), connectivity (60, 61), and function (60, 62). Several of such “precision neuroimaging” datasets have already led to an advanced characterization of functional systems (63, 64) or fostered enhanced microstructural modelling, but previous precision imaging datasets were either prioritizing functional or structural imaging acquisitions, and rarely both in the same subjects. Moreover, prior precision imaging investigations were mainly carried out at 3T. In this study, we expand this work by leveraging a recently introduced precision neuroimaging (PNI) dataset, which combines repeated high-resolution structural and functional acquisitions at 7T, offering an opportunity to interrogate cortical organization in the living human brain with high sensitivity and specificity.

The current work examined the interplay of local cortical arealization and global integration. Leveraging probabilistic cytoarchitectonic maps of the Julich-Brain atlas (16), we subdivided the cortex into 228 areas. In those, we profiled microstructural, structural, and functional gradients derived from repeated 7T MRI scans. We then examined how multimodal gradient profiles differed across areas. As local-global cortical organization is presumably tied to cognitive functional architecture, we cross-referenced our maps to multiple fMRI tasks conducted in the same participants, and in particular studied the relation between inter-area gradient profiles and functional diversity across different tasks acquired in the same subjects. By integrating measures of cortical cytoarchitecture with multimodal high-definition MRI, our work sheds light on local-global cortical organization and advances our understanding of cortical structure-function relationships.

## Results

### Bridging local and global cortical organization

We constructed high-resolution cortex-wide connectomes, encompassing microstructural profile covariance (MPC) (20), structural connectivity (SC) (65), and functional connectivity (FC) (19, 66), in 10 healthy adults who underwent three repeated multimodal MRI scans at 7T (**Figure 1A**). We estimated connectome eigenvectors that characterized spatial gradients of MPC, SC, and FC. We focused on the first five gradients across modalities (MPC: 31%; SC: 18%; FC: 25%). In line with prior work (20), the principal MPC gradient was anchored on one end by primary sensory areas and on the other end by paralimbic regions. The principal SC gradient exhibited anterior-posterior axis, clearly dividing the cortex into two parts bounded by sensorimotor areas, as reported previously (67). The first FC gradient differentiated sensorimotor cortices and default mode network, recapitulating earlier work (19, 66). Other gradients were also in keeping with prior reports (**Figure 1A**) (20, 65-67). Finally, gradients were averaged in areas derived from the Julich-Brain atlas (**Figure 1B**), to generate an area-wise gradient profile matrix.

**Figure 1.**
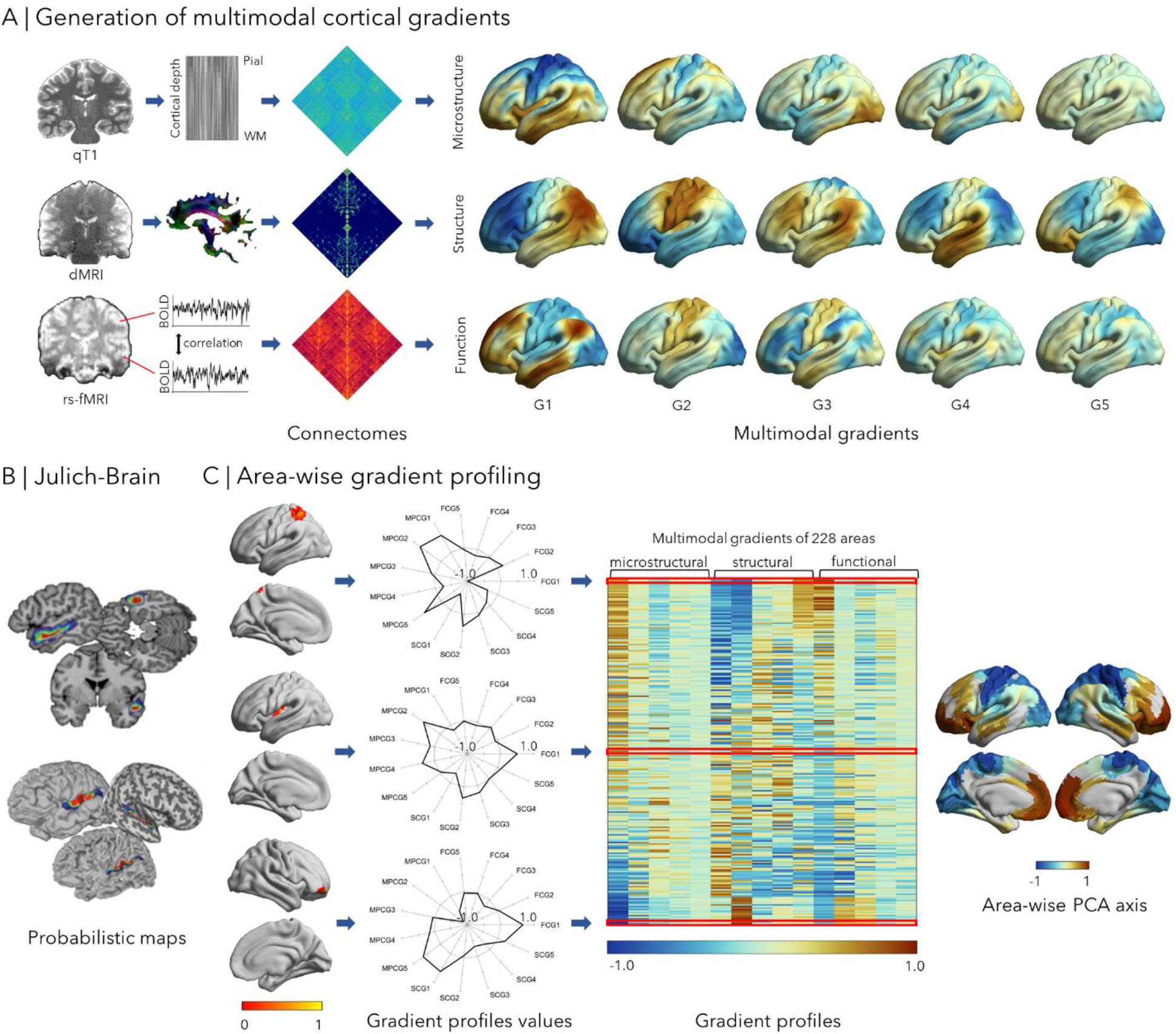
Integration of global cortical gradients with cortical arealization. **(A) Generation of multimodal cortical gradients.** Cortex-wide connectomes were constructed from microstructural profile covariance (MPC) (20), structural connectivity (SC) (65, 67), and functional connectivity (FC) (19, 66) at a vertex-level based on repeated 7T MRI. We applied non-linear dimensionality reduction techniques (68) to each connectome and aggregated the first five eigenvectors/gradients. **(B) Cortical arealization: Julich-Brain Atlas**. Probabilistic area definitions were derived from the Julich-Brain atlas (16), a *post-mortem* cytoarchitectonic atlas based on the mapping of areas ten postmortem brains, and their superimposition in MNI space **(C) Area-wise gradient profiling**. We averaged vertex-wise gradients in each of the 228 areas, producing area-specific multimodal gradient profiles. *Left panel:* the resulting gradient profiles located in the middle and at the two ends of the main axis were visualized in spider plots. *Middle panel:* the multimodal gradient profiles. *Right panel:* the first principal component from the multimodal gradient profiles.

### Cortical patterns of local-global integration

We examined the similarity and differences of gradient profiles across areas. To this end, we first conducted principal component analysis (PCA) on the multimodal gradient profiles. This multimodal gradient arranges cortical areas in terms of the similarity or their gradient values and followed a sensory-fugal axis in multimodal gradient profiles, anchored on prefrontal/cingulate regions on the one end and central/occipital regions on the other end, integrating salient features of its constituents (*i*.*e*., the individual MPC, SC, and FC gradients) in a synoptic manner (**Figure 1C**).

To further quantify area-to-area differences, we computed an inter-area cosine distance matrix (**Figure 2A**). The mean value of each row in this matrix indicates the overall dissimilarity of a given area from all other areas in terms of the multimodal gradient profiles. To identify cortical areas with significantly higher/lower dissimilarity compared to all other areas, we conducted spatial permutation tests (1000 permutations) that randomly rotated the Julich-Brain atlas on a sphere (69). We found significant and highest global dissimilarity in areas of the sensorimotor cortex (p_spin_<0.05, false discovery rate (FDR) correction; **Figure 2A**), indicating that these areas are the most unique across the cortex in terms of their multidimensional gradient profiles. Conversely, we observed the lowest dissimilarity in insular and fronto-temporal regions after spatial permutation tests (p_spin_<0.05, FDR correction).

**Figure 2.**
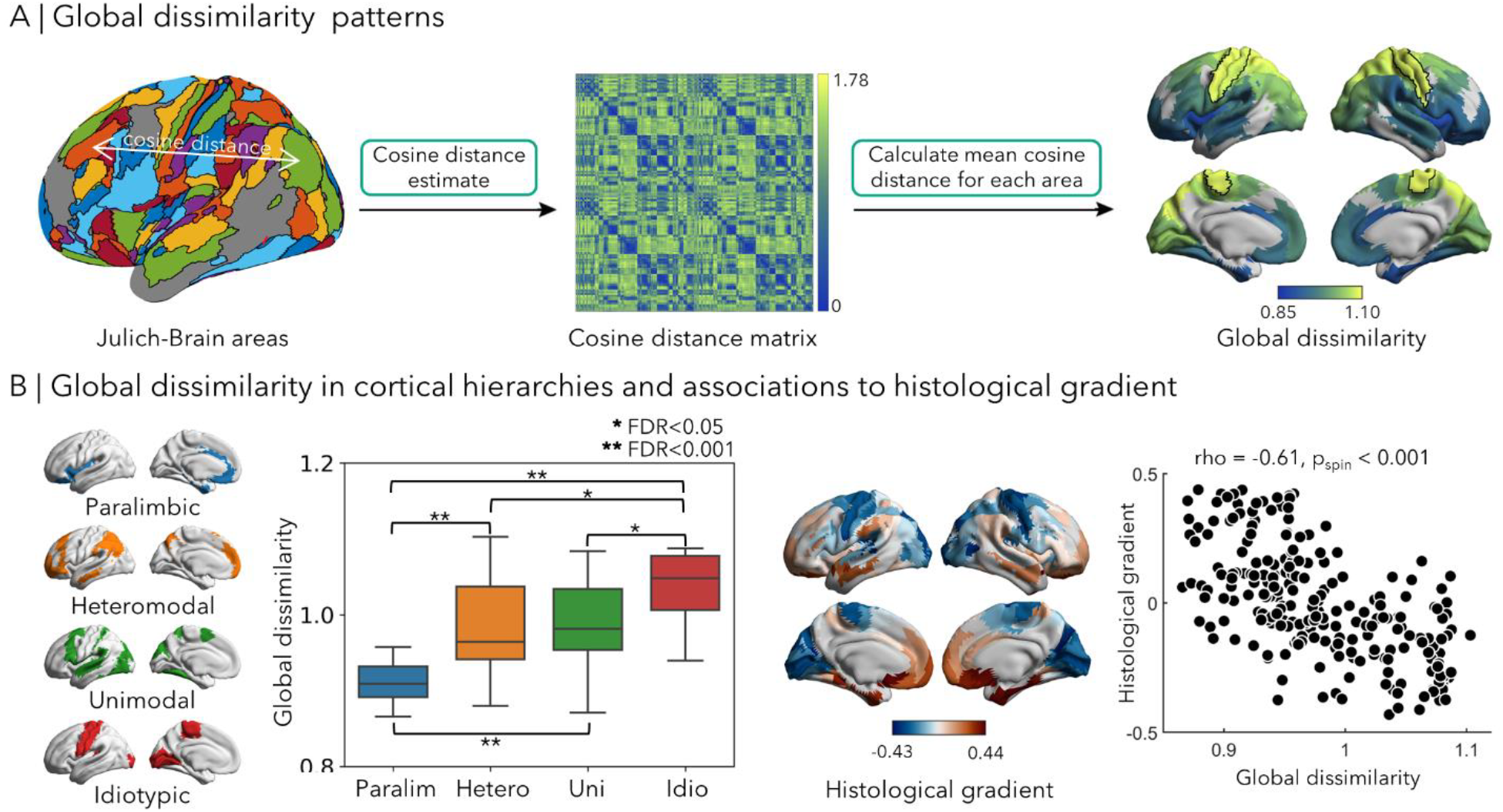
Inter-area heterogeneity. **(A) Global dissimilarity patterns**. We calculated the cosine distance between each pair of cortical areas and computed the mean value for each area. To identify cortical areas with the highest and lowest global dissimilarity, we conducted 1000 permutation tests (69). Regions with significantly higher global dissimilarity compared to other areas after applying FDR correction were highlighted using black boundaries. **(B) Global dissimilarity in cortical hierarchies and associations to histological gradient**. The *left panel* illustrates the distribution of global dissimilarity across four cortical hierarchies. To examine differences between each cortical hierarchical level, two-sample t-tests were conducted with FDR correction. To explore associations with the histological gradient, Spearman’s correlation coefficients were computed, and p-values corrected using spin permutation tests (69).

To gain a deeper understanding of these patterns, we investigated the distribution of global dissimilarity across four cortical hierarchical levels derived from a prior taxonomy of the primate brain proposed by Mesulam (70). Using a two-sample t-test, we compared global dissimilarity between each pair of cortical hierarchical levels (i.e., paralimbic, heteromodal, unimodal and idiotypic). The results revealed that the idiotypic system had the highest global dissimilarity compared to other systems (p_spin_<0.05, FDR correction, **Figure 2B**). In contrast, paralimbic systems showed lowest global dissimilarity (p_spin_<0.05, FDR correction), aligning with prior findings. To further explore associations between global dissimilarity and cortical microstructure, we generated a MPC matrix of histological data obtained from the BigBrain dataset (71), a 3D reconstruction of *post-mortem* human brain histology, and estimated its principal gradient. This gradient has previously been shown to closely recapitulate Mesulam’s taxonomy of cortical hierarchical organization. In effect, we also observed a significant correlation between the histological gradient and global dissimilarity (rho=-0.61, p_spin_<0.001).

To examine cortical similarities using an alternative approach, we also performed hierarchical clustering on cortical similarity matrices and found similar results, providing four robust clusters recapitulating sensory-fugal hierarchies (**Figure S1, Supplementary Materials**). Moreover, we investigated intra-area heterogeneity at the vertex level. Here, we calculated the cosine distance between the gradient profile of each cortical vertex and the mean gradient profile of the area to which it belongs (**Figure S2A**). We found that intra-area heterogeneity was considerably lower compared to inter-area heterogeneity. Comparing hierarchical levels, we did not observe the same relationship as for inter-area heterogeneity. In particular, intra-area heterogeneity was higher in heteromodal and unimodal association systems, compared to idiotypic and paralimbic regions (p_spin_<0.05, FDR correction) there was no significant association to the histological gradient derived from BigBrain (rho=-0.17, p_spin_=0.12; see **Figure S2** and **Supplementary Materials**).

### Associations to functional diversity across task states

The cortical layout is intricately linked to cognition (20, 72). To investigate how functional connectivity patterns change across diverse cognitive states, we administered nine different fMRI tasks, including episodic memory encoding and retrieval, pattern separation encoding and retrieval, semantic retrieval, and four passive movie watching paradigms in the same participants at 7T. We constructed functional connectivity for all states and calculated the cosine distance between the corresponding whole brain functional connectivity matrices to estimate cross-state diversity for each area (**Figure 3A**). We observed highest functional variability in the medial temporal lobe and orbitofrontal cortex, while the medial frontal lobe and primary sensory cortex had low variability (**Figure 3B**). To further explore how this functional variability relates to cortical organization, we assessed associations between the cross-state diversity map and inter-area dissimilarity. This was done by computing Spearman’s correlation coefficient and correcting p-values using 1,000 spin permutation tests. Notably, we identified a marked correlation between cross-state diversity and global dissimilarity (rho=-0.71, p_spin_<0.001; **Figure 3B**). To control for potential influences from tSNR, we performed partial correlation analysis with tSNR as a covariate and found consistent results (p_spin_<0.001). Again, we only found weak, and marginally significant association with local dissimilarity (rho=-0.19, p_spin_=0.065; **Figure S2**).

**Figure 3.**
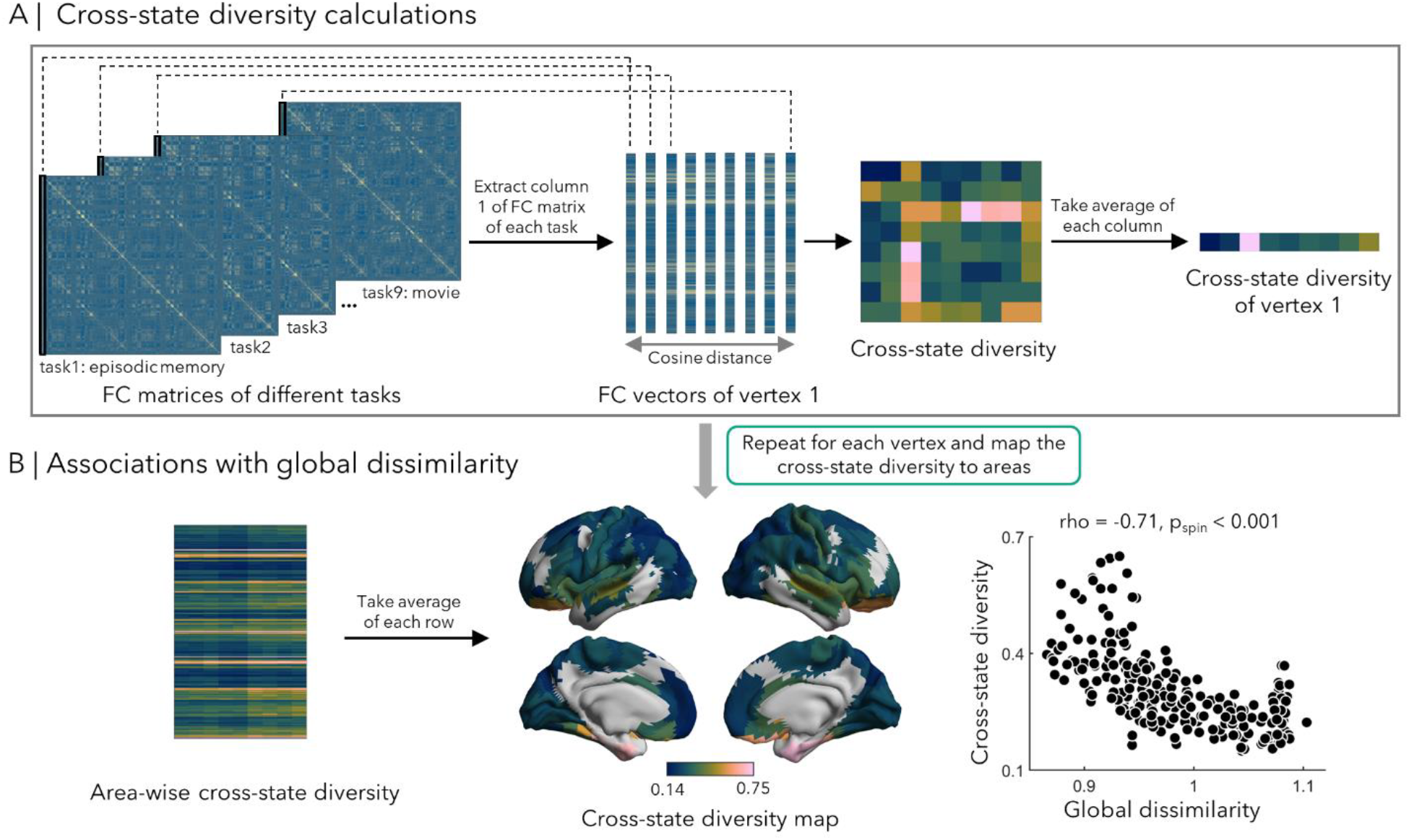
Associations to cross-state functional diversity. **(A) Cross-state diversity calculations**. The FC matrices were generated using time series data obtained from for nine tasks fMRI sessions in the same subjects. This was followed by the estimation of cosine distances across different tasks. This process resulted in a cross-state diversity matrix of dimensions. To quantify the cross-state diversity for the first vertex of all tasks, the average of each column in this matrix was computed. **(B) Associations with global dissimilarity**. By repeating the procedures outlined in *Panel A* for each vertex and subsequently mapping the results to areas, we created an area-wise cross-state diversity map. Spearman’s correlation coefficient was computed, and p-values were corrected using spin permutation tests.

### Robustness with respect to analysis parameters

To assess robustness of our findings with respect to analysis parameters, we recalculated gradients, gradient profiles, global and local dissimilarity for each modality using different thresholds of the connectivity matrix (50%, 60%, 70%, 80%, 90%). Correlations between global and local dissimilarity derived from gradient profiles with different thresholds were assessed, revealing consistent results (**Figure S3**). Moreover, we investigated the impact of varying the number of gradients within each modality, ranging from three to seven. Global and local dissimilarity was estimated, and consistent results were observed across different numbers (**Figure S3**).

### Reliability at the single subject level

We assessed our findings at each of the 10 individual participants that were scanned at 7T. Similar results were found across participants, including gradient profile matrices and global dissimilarity (**Figure 4A**). Moreover, we observed marked negative correlations between global dissimilarity and the histological gradient (rho=-0.58±0.047, ranged from -0.63 to -0.49, all p_spin_<0.001), and between global dissimilarity and cross-state diversity (rho=-0.53±0.067, ranged from -0.64 to - 0.45, all p_spin_<0.001). Global dissimilarity across subjects were highly correlated (r=0.90±0.008; **Figure 4B**).

**Figure 4.**
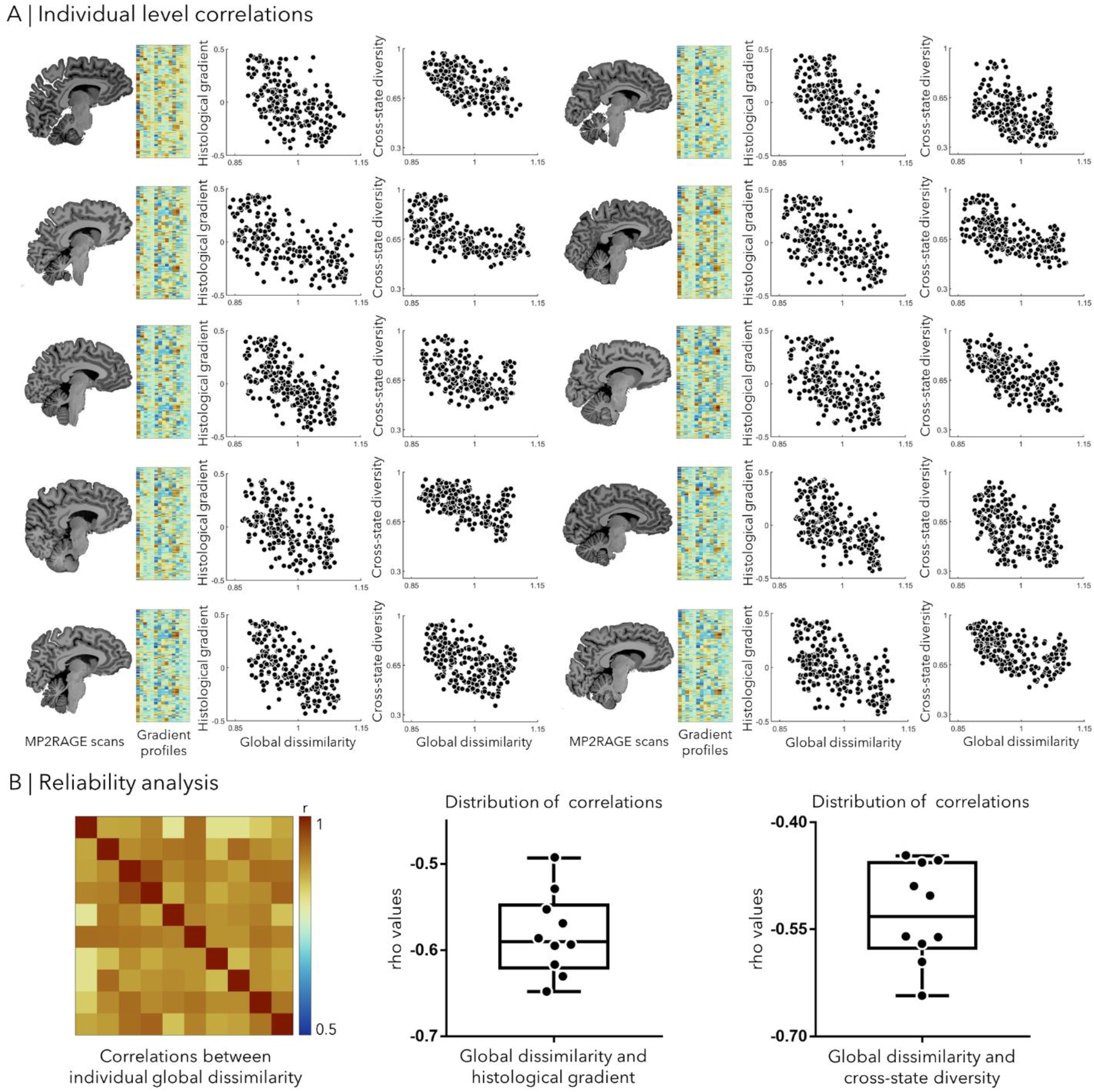
Reliability analysis at the single subject level. **(A) Individual level correlations**. For each of the 10 subjects, we generated gradient profiles and global dissimilarity. Associations between individual global dissimilarity and histological gradient, as well as cross-state diversity were examined using Spearman’s correlation coefficients. **(B) Reliability analysis**. Associations between individual global dissimilarity were estimated using Pearson’s correlation coefficient. Distributions of rho values between global dissimilarity and histological gradient (mean±SD of rho values=-0.58±0.047), and with cross-state diversity were examined (mean±SD of rho values=-0.53±0.067). Abbreviation: MP2RAGE, magnetization-prepared 2 rapid gradient echo.

### Replication at 3T

To ensure the reproducibility of our findings, we conducted a replication analysis involving an independent sample of 50 healthy young adults (age: 29.54±5.62 years, 23 females) scanned at 3T (67). Results from the replication sample revealed consistent findings, with similar multimodal gradients, gradient profiles, and inter-area heterogeneity (**Figure S4A-B**). Notably, associations with histological gradients (rho=-0.41, p_spin_<0.001) and cross-state diversity (rho=-0.62, p_spin_<0.001) remained consistent. Again, we observed only a marginal association between local dissimilarity and cross-state diversity (rho=-0.11, p_spin_>0.1; **Figure S4C**), with no significant correlation found for histological gradients.

## DISCUSSION

Functional specialization and integration are two cornerstones of neural organization (73, 74). While specialization relates to distinctive neural behavior across different contexts (75), functional integration emphasizes the shared influence among regions, ultimately contributing to coherent experiences and behavior (75, 76). The present study combined multimodal MRI acquisitions with gold standard descriptions of cortical cytoarchitecture (16), in order to identify the similarity and dissociation of global cortical fingerprints across areas. Vertex-wise multimodal connectomes were constructed from high-field 7T MRI data, and cortical gradients estimated, aligning with those described in prior studies (19, 20, 65). We noted higher global dissimilarity in sensorimotor cortices and lower global dissimilarity in the transmodal system, indicating the distinctiveness of the primary sensory cortex which supported its functional specialization. Significant associations between cross-state functional variability and inter-area heterogeneity were identified, indicating a link between global cortical motifs and functional diversity across different task contexts. These findings suggest a sensory-paralimbic differentiation in cortical gradient fingerprints, providing insights into neural motifs contributing to specialized and integrative brain functions.

The availability of multimodal neuroimaging data offers opportunities for examining brain organization across different spatial scales (36, 37, 77). However, investigations into principles governing functional segregation *versus* specialization are often constrained by the resolution of *in-vivo* MRI (76). To address this limitation, our study leveraged a high-field *in-vivo* 7T MRI dataset that features high signal strength, high spatial resolution, and the availability of several modalities (78). Data were acquired across multiple time points to further improve MR signal-to-noise ratio and contrast-to-noise ratio, thereby enhancing reliability and robustness (60-64). To integrate descriptions of segregation and integration, we utilized a gold standard atlas derived from *post-mortem* histological data to finely partition the brain into distinct regions based on cytoarchitecture (16) and combined this with dimensional gradient descriptions that situate each cortical area into the larger tapestry of cortical microstructure, connectivity, and function (19). Given that the gradient profiles in our study represent vectors in a multidimensional space, we then measured the distance between all cortical areas using a cosine distance metric. Interestingly, we observed an overarching pattern of cortical global dissimilarity, with one end featuring sensory and motor cortices featuring the highest global dissimilarity and the other end encompassing heteromodal and paralimbic areas in the transmodal apex. This suggests that functional specialization in primary sensorimotor areas is accompanied by a more distinctive organization pattern. Overall, the axis of global dissimilarity is consistent with the gradients previously reported for single modalities that are, especially at the level of function and microstructure, also anchored in primary systems on the one end, and more heteromodal and paralimbic regions on the other end (19, 20). Tract-tracing and neuroimaging experiments have documented that primary sensory and motor cortices host more short-range cortico-cortical connections than transmodal systems (66), and these regions also tend to have a higher coupling between microstructure and function, and between structural connectivity and function (30). Such findings are potentially in support of their more specialized functional profiles (66, 79). Notably, the pattern of cortical organization within the limbic system closely aligns with that of other cortical areas, extending previous findings derived from single modalities (19, 20, 79, 80). This supports the integrative role of the limbic system, which may enable it to participate in various cognitive processes (81).

Distance-dependence theory suggests that regions in close proximity are more likely to exhibit inter-connections (82, 83). Moreover, cross-species research indicates that areas sharing similar microstructural and neurobiological characteristics tend to be interconnected (84-87). These findings support the overarching idea that adjacent regions with short-range connections often share gene expression and microstructural similarities, contributing to their specialized functional roles. Conversely, while nearby neurons are expected to share similar microstructural properties when extending smooth macroscale topography to the microscale (88), prior electrophysiological experiments in various mammalian brain regions have shown that nearby neurons can exhibit disparate response properties (89-93). Thus, our study examined the intra-area heterogeneity of brain organization to probe whether individual functional units (i.e. populations of neurons within a voxel) are more consistent or more unique from their neighbours. Consistent with our expectations, overall intra -area variability was relatively low when compared with variations between areas, confirming the utility of parcellations as a concept reflected in most atlases, more generally.

We also observed notable subregional variations within areas, and these were most pronounced in uni-as well as heteromodal association cortices, in line with their more integrative functional role, compared to primary sensory and motor areas. Contrary to these inter-area findings, we did not observe significant trends or associations at the level of intra-area variations to the functional and microstructural measures of cortical hierarchy investigated in the study. By nevertheless showing an overall increased intra-area heterogeneity in association cortices, our findings may support the vital role of association cortex in information integration (7). We speculate that intraregional heterogeneity can support the processing and integration of inputs across a large cortical territory (94). That is, for disparate types of information to be integrated, they must at minimum be present in the same brain regions. Moreover, these findings are also aligned with the tethering hypothesis of cortical patterning, where a disproportionate enlargement of uni- and heteromodal systems during human evolution and the progressive decrease of genetically mediated signalling gradients may have contributed to their higher local dissimilarity and relative structure-function decoupling (27, 95).

To tie our findings to cortex-wide functional differences, we assessed spatial correlations between inter-area dissimilarity as well as functional diversity across different fMRI tasks obtained in the same subjects. The observed negative correlation between global dissimilarity and cross-state diversity suggests that globally more specialized regions, such as sensorimotor and visual cortices, also exhibit more stable functional connectivity patterns across various tasks. A recent study reported that functional connectivity in somatomotor cortex increased with age during childhood through adolescence, whereas it declined in association cortices, reinforcing the differentiation of sensorimotor and association systems in typical development (96). These findings support the existence of a sensorimotor-association axis of cortical organization (8, 19), and may explain the observed higher stability of functional connectivity in sensorimotor cortex across tasks. On the other hand, paralimbic areas presented with least distinctiveness compared to other cortical regions, and also showed the highest cross-state diversity in functional connectivity. This may be attributed to the involvement of these regions across a diverse array of cognitive and affective processes (81, 97), and prior findings suggesting overall increased signalling variability in these systems (98). Collectively, these findings suggest that the heterogeneity in global cortical motifs across different regions is also reflected in their diverse participation across different functional contexts.

A series of robustness analyses, exploring the influence of thresholds for gradient estimation and the number of gradients, yielded similar results, suggesting that our analyses was not affected by variations in specific analysis parameters. Moreover, we observed consistent findings at the level of individual participants and could replicate our findings using a completely independent dataset scanned at 3T. It is nevertheless essential to note that there may be additional sources of inter-participant variability and cross-scale effects that could be further explored in future studies (99). Notably, the probabilistic atlas utilized in this study to define cortical areas did not cover the entire cortex; some cortical areas remain undefined, presenting a limitation. However, this limitation is anticipated to be resolved with the publication of the whole-brain probabilistic map in the future, as it is currently under development (16). As our work shows, cortical parcellation and gradients both can provide synergistic information to integrate histological and neuroimaging data in the human brain. By thus reconciling local and global cortical features, our work may provide new insights into the neuroanatomical basis of specialized and integrative cortical functions.

## Methods

### Participants

Our study used two independent human neuroimaging datasets. A 7T dataset (MICA-PNI, 10 subjects, multiple time points) was used for discovery and reliability assessment. A 3T dataset (MICA-MICs, 50 subjects, one time point) was used for replication.

a. *MICA-PNI*. For our main analysis, we investigated the imaging and phenotypic data of 10 unrelated healthy adults (age: 29.20±5.20 years, 5 females). Each participant underwent three sessions on separate days. Data were collected between March 2022 and June 2023. This dataset is openly available at the OSF platform (https://osf.io/mhq3f/).
b. *MICA-MICs* (67). This dataset consisted of 50 unrelated healthy young adults (age: 29.54±5.62 years, 23 females). Data were collected between April 2018 and February 2021. This dataset has recently been released and is openly accessible to the imaging community (https://portal.conp.ca/dataset?id=projects/mica-mics) (67).

Both studies were approved by the Ethics Committee of the Montreal Neurological Institute and Hospital, and written and informed consent were obtained from all participants.

### MRI acquisition

#### MICA-PNI

Scans were acquired using a 7T Siemens MAGNETOM Terra scanner equipped with an 8/32-channel transmit/receive Nova head coil at the McConnell Brain Imaging Centre (BIC) of the Montreal Neurological Institute. Each participant underwent multiple types of scans, including magnetization-prepared 2 rapid acquisition by gradient echo (MP2RAGE), multi-shell diffusion-weighted imaging (DWI), rs-fMRI, and task fMRI.

MP2RAGE is a self bias-field corrected sequence that acquires two images with different inversion times to generate a myelin-sensitive map of the T1 relaxation times of the brain and a synthetic T1-weighted (T1w) image. The 3D MP2RAGE sequence parameters are the following: 0.5mm isotropic voxels, matrix=320×320, 320 sagittal slices, repetition time (TR)=5170 ms, echo time (TE)=2.44 ms, inversion time (TI)=900 ms, flip angle=4°, iPAT=3, TI1=1000 ms, TI2=3200 ms, bandwidth=210 Hz/px, echo spacing=7.8 ms, and partial Fourier=6/8. Scans were visually inspected to ensure minimal head motion, and rescanned if necessary. Both inversion images were combined for T1 mapping to minimize sensitivity to B1 inhomogeneities and optimize intra- and inter-subject reliability (100, 101).

DWI data was acquired using a multiband accelerated 2D spin-echo echo-planar imaging sequence. The acquisition included three shells with b-values of 300, 700, and 2000 s/mm2, and 10, 40, and 90 diffusion weighting directions per shell, respectively. The parameters used were: 1.1 mm isotropic voxels, TR=7383 ms, TE=70.60 ms, flip angle=90°, refocusing flip angle=180°, FOV=224×224 mm^2^, slice thickness=1.1 mm, multi-band factor=2. Reverse phase encoding b0 images were obtained for distortion correction of the DWI scans.

For rs-fMRI, a 6-minute scan was conducted using multiband accelerated 2D-BOLD gradient echo-planar imaging. The parameters for rs-fMRI were as follows: 1.9 mm isotropic voxels, TR=1690 ms, TE1=10.8 ms, TE2=27.3 ms, TE3=43.8 ms, flip angle=67°, FOV=224×224 mm2, multiband factor=3, and echo spacing=0.53 ms. Participants were instructed to keep their eyes open, avoid falling asleep, and fixate on a cross presented on the screen. Two spin-echo images with opposite phase encoding directions were also acquired for distortion correction of the rs-fMRI scans, with the following parameters: phase encoding=AP/PA, 1.9 mm isotropic voxels, FOV=224×224 mm^2^, slice thickness=1.9 mm, TR=3000 ms, TE=18.4 ms, flip angle=90°. Using similar parameters, we also collected multiple task fMRI scans in three sessions, including episodic memory, semantic memory, spatial memory, MST and four different movie tasks.

#### MICA-MICs

All scans were acquired using a 3T Siemens Magnetom Prisma-Fit equipped with a 64-channel head coil at the McConnell Brain Imaging Centre of the Montreal Neurological Institute. All participants underwent T1w structural MRI, DWI, multiband rs-fMRI, and qT1 imaging. For details of the parameters of sequences please see the **Supplementary Materials**.

### Multimodal MRI processing

Raw DICOMS were sorted, and converted to NIfTI format using dcm2niix (102). The folder structure was verified using the BIDS (103) validator (https://bids-standard.github.io/bids-validator/). All pre-processing steps described below were implemented using open-source pipeline micapipe v0.2.3 available at (http://github.com/MICA-MNI/micapipe) (104).

The MP2RAGE scans of each subject were reoriented using FSL (105), linearly co-registered, averaged, with background noise removed, corrected for intensity nonuniformity using N4 bias field correction from ANTS (106), and segmented into white and grey matter using FSL FAST (105). Resulting volumes were skull stripped using FSL (105, 107). Cortical surface models were generated from native T1w scans using FastSurfer (108). Surface reconstructions for each subject underwent manual correction for segmentation errors, by placing control points and applying manual edits.

Regarding the DWI data, pre-processing was carried out using MRtrix (109) in the native DWI space. The DWI data underwent denoising (110, 111), b0 intensity normalization, and correction for susceptibility distortion, head motion, and eddy currents. These corrections were performed using FSL (112) and involved utilizing two b=0 s/mm^2^ volumes with reverse phase encoding. Anatomical masks for tractography were non-linearly co-registered to native DWI space using the deformable SyN approach implemented in ANTs (113).

For the rs-fMRI scans, pre-processing steps were conducted using AFNI (114) and FSL(105) tools. The first five volumes were discarded to ensure magnetic field saturation. We applied Multi-Echo Independent Components Analysis (ME-ICA) (115, 116) to improve the signal-to-noise ratio and effect of motion correction. Spike regression was applied to remove timepoints with large motion spikes, effectively removing nuisance signals (117, 118). The volume timeseries were registered to FastSurfer (108) space using boundary-based registration implemented in ANTs using linear and non-linear methods (119).

For details of the MRI processing of MICA-MICs dataset, please see the **Supplementary Materials**.

### Generating multimodal connectome matrices

To investigate the vertex-wise multimodal connectomes, we first constructed a downsampled fsLR-5k surface using HCP’s workbench tools (wb_command) (120). The fsLR-32k surface templates and resampling spheres between ‘fsaverage’ and ‘fs_LR’ were accessed from the HCP’s open-access pipeline (121). Subsequently, we downsampled the surface template, registration spheres, and mid-wall mask to 5k, resulting in a mesh comprising 4432 cortical vertices for each hemisphere. All vertex-wise analyses were performed based on this fsLR-5k surface.

We calculated vertex-wise MPC matrices for each participant. Consistent with previous work (20, 122, 123), we constructed 14 equivolumetric surfaces between the pial and white matter boundaries to sample qT1 intensities across cortical depths. This procedure generated distinct intensity profiles reflecting intracortical microstructural composition at each cortical vertex. Data sampled from surfaces closest to the pial and white matter boundaries were removed to mitigate partial volume effects. Intensity values at each depth were mapped to a common template surface, resampled to fsLR-5k surface, and spatially smoothed across each surface independently (full width at half maximum [FWHM]=3mm). Vertex-wise intensity profiles were cross-correlated using partial correlations controlling for the average cortex-wide intensity profile, and log-transformed. This procedure resulted in the MPC matrices representing participant-specific similarity in myelin proxies across the cortex.

To generate each individual’s SC, we employed MRtrix on pre-processed DWI data (109). Each surface vertex from the fsLR-5k surface was translated into a volumetric region of interest that filled the cortical ribbon using workbench tools(120). This process yielded approximately 10k seeds/targets for structural connectome generation. Anatomical segmentations and volumetric seeds were then mapped to DWI space applying the non-linear registration warp-field mentioned earlier. Next, we estimated multi-shell and multi-tissue response functions (124) and performed constrained spherical deconvolution to derive a fiber orientation distribution map (125, 126). This procedure, achieved through MRtrix, generated a tractogram with 40M streamlines, with a maximum tract length of 250mm and a fractional anisotropy cutoff of 0.06. To reconstruct whole-brain streamlines weighted by cross-sectional multipliers (127), we applied spherical deconvolution informed filtering of tractograms (SIFT2). Connection weights between seeds/targets were defined as the streamline count after SIFT2.

Next, individual rs-fMRI timeseries mapped to subject-specific surface models were resampled to fsLR-5k surface. Surface-based rf-MRI data underwent spatial smoothing with a Gaussian kernel (FWHM=3mm). An individual’s FC matrix was generated by cross-correlating all vertex-wise timeseries. Correlation values subsequently underwent Fisher-R-to-Z transformations. FC matrices of all task fMRI scans were also generated using the same approach.

### Construction of gradient profiles

We converted each participant’s SC, MPC, and FC matrices to a normalized angle affinity matrix respectively, and applied diffusion map embedding on these matrices to generate multimodal gradients (128). This non-linear dimensionality reduction procedure identified eigenvectors that describe main spatial axes of variance. Procrustes analysis aligned subject-level gradients to a group-level template generated from the group-average matrix of all participants. Gradients of the right hemisphere were aligned to the left hemisphere. Gradient analyses were performed using BrainSpace (v0.1.10; http://github.com/MICA-MNI/BrainSpace), limiting the number of gradients to 10 and using default sparsity (keeping the top 10% of SC weights) and diffusion (α=0.5) parameters (68). Here, we focused on the first five principal gradients of each modality (**Figure 1A**). For each modality, all gradients were normalized by dividing by the maximum value in the absolute value of gradients, with values ranging from -1.0 to 1.0.

The Julich-Brain is a 3D probabilistic atlas of the human brain’s cytoarchitecture, resulting from the analysis of 10 *post mortem* human brains (16) (**Figure 1B**). The probabilistic cytoarchitectonic maps (Julich-Brain v2.9, https://julich-brain-atlas.de/) were projected onto a template fsLR-5k surface to generate a surface-based representation (16). Surface-based probabilistic maps contained values indicating the probability of an area being localized in each voxel, ranging from 0% to 100% overlap, with values ranging from 0 to 1. We registered the probabilistic maps to the fsLR-5k surface template. For each vertex, we defined its area label by identifying the area with the highest probability at that position. This area label was then used to assign all vertices on the fsLR-5k surface to the 228 areas defined by the Julich-Brain.

In accordance with the previous sections, we computed the first five gradients of microstructure, structural connectivity, and functional connectivity, which accounted for the 31%, 18%, and 25% of variance in the input data (19). This allowed us to assign a unique gradient profile to each vertex, consisting of 15 gradient values (three modalities multiplied by five gradients). Subsequently, we employed the probabilistic map of the Julich-Brain to generate area labels and assign cortical vertices to cytoarchitecturally-defined regions. Consequently, we obtained area-wise gradient profiles that effectively captured the majority of variance present in multimodal connectomes. To investigate the hierarchy of gradient profiles, we conducted PCA on the gradient profiles, aiming to identify the primary axis across all areas. We then arranged the area-wise gradient profiles based on the first component of the PCA and examined the gradient profile patterns of the areas located in the middle and at the two ends of the derived PCA axis.

### Gradient profile analyses

#### Inter-area heterogeneity and homogeneity assessment

Understanding the relationships between diverse brain regions, including their similarities and differences, is essential for investigating the spatial patterns of brain organization. In this study, we focus on exploring inter-area heterogeneity and homogeneity to further reveal the global layout of cortical area. To quantify inter-area heterogeneity, we computed the global dissimilarity for each area. Specifically, we calculated the cosine distance between gradient profiles of each area, resulting in a cosine distance matrix (**Figure 2A**). The global dissimilarity was defined as the mean of each row in the cosine distance matrix, representing the distance between an area and all other areas. To identify cortical regions with significantly higher or lower global dissimilarity, we projected the global dissimilarity map onto a sphere and conducted 1,000 spin permutation tests. An area was considered to have the highest global dissimilarity among the cortex if its original global dissimilarity value exceeded 97.5% of the permutation values. Conversely, an area was regarded as having the lowest global dissimilarity if its original value was lower than 97.5% of the permutation values. To correct for multiple comparisons, we applied the FDR correction. To further investigate patterns of global dissimilarity, we conducted a network-level analysis utilizing the scheme proposed by Mesulam (70), which delineates four cortical functional zones (i.e., idiotypic, paralimbic, unimodal, heteromodal; see **Figure 2B**). Global dissimilarity between each pair of cortical hierarchies was compared using a two-sample t-test. FDR corrections were applied to correct for multiple comparisons, while spatial autocorrelation spin permutation tests were conducted for all tests. In order to explore the associations between global dissimilarity and cortical microstructural hierarchy, we generated a MPC matrix based on histological data from BigBrain (71), an ultrahigh-resolution 3D human brain model. From this matrix, we estimated the principal histological gradient as a representation of microstructural hierarchy. To examine the associations between the histological gradient and global dissimilarity, we calculated Spearman’s correlation coefficient, with p-values corrected using 1,000 spin permutation tests.

Differences between cortical regions are crucial to functional specialization, but at the same time, similarities between regions support the realization of higher-order cognitive functions and functional integration across brain regions. To assess the homogeneity of cortical areas, we calculated (1 - cosine distance) to represent the similarity between regional gradient profiles, resulting in an affinity matrix (**Figure S1A**). To evaluate the association between inter-area similarity and cortical hierarchy, we examined the distribution of similarity coefficients across four hierarchy levels. To identify cortical area with higher similarity, we performed hierarchical clustering on the affinity matrix to detect groups among the areas. We evaluated the clustering performance by calculating criterion values to determine the optimal number of clusters. We scrutinized the clusters with the highest criterion value and assessed the distribution of cortical hierarchies within each cluster to investigate the association between inter-area homogeneity and cortical laminar differentiation.

#### Intra-area homogeneity and heterogeneity assessment

In addition to the relationships between cortical areas, exploring the layout within an area will provide insights into understanding the local organization of the cortex. Given that each area was originally defined based on shared neuroanatomical features, we expect to find overall high intra-area homogeneity. Here we quantified, however, to which extent the level of homogeneity varies across the brain. We assessed intra-area heterogeneity at both the region-level and network-level. As previously described, we calculated the gradient profiles for each area by averaging the vertex-wise gradients within that area. For a given area *i*, we calculated the cosine distance between the vertex-wise multimodal gradients and gradient profile of area *i*, resulting in the generation of local dissimilarity of area *i* (**Figure S2**). This procedure was repeated for all areas, yielding the vertex-wise local dissimilarity map.

To visualize the patterns of intra-area heterogeneity more effectively, we obtained the average of the vertex-wise local dissimilarity within each area. Furthermore, we investigated the local dissimilarity at the network-level using the four cortical hierarchies proposed in a previous study (70). We examined the distribution of local dissimilarity within these four cortical hierarchies and compared the differences between each hierarchy using a two-sample t-test with FDR correction and spatial autocorrelation spin permutation tests. To investigate associations between local dissimilarity and cortical microstructural hierarchy, we calculated Spearman’s correlation coefficient between the histological gradient and local dissimilarity, correcting the p-value using 1,000 spin permutation tests.

#### Associations to cross-state functional variability

To explore patterns of functional variability across different task fMRI scans, we constructed FC matrices using timeseries data derived from multiple task fMRI sessions. The cosine distance was then computed for each vertex across different tasks, resulting in a cross-state diversity matrix (**Figure 3A**). To quantify the cross-state diversity for a specific task, we averaged all distance values between that task and others. This process was repeated for all tasks, and the outcomes were mapped to areas, generating area-wise cross-state diversity for each task. Focusing on overall variability, we calculated the average of values across all tasks to create the cross-state diversity map. To investigate associations between cross-state diversity and global/local dissimilarity, we computed Spearman’s correlation coefficient. The resulting p-values were corrected using 1,000 spin permutation tests.

## Data availability

The 7T MRI data is openly available at the OSF platform (https://osf.io/mhq3f/) upon publication. The Julich-Brain atlas is available at https://julich-brain-atlas.de/ and provided through the EBRAINS platform (https://www.ebrains.eu/tools/human-brain-atlas). The MICA-MICs replication data is openly available at https://portal.conp.ca/dataset?id=projects/mica-mics (67). Gradient mapping analyses was based on BrainSpace (https://brainspace.readthedocs.io/en/latest/) (68).

## Code availability

Code for MRI data preprocessing is available at https://github.com/MICA-MNI/micapipe (104). The code for connectome gradients generation is available at https://github.com/MICA-MNI/BrainSpace. Code for the main analysis is openly available on https://github.com/MICA-MNI/micaopen/tree/master/gradient_profiles.

## Acknowledgements

Yezhou Wang, Dr. Alan Evans, and Dr. Boris Bernhardt were supported by the Helmholtz International BigBrain Analytics and Learning Laboratory (HiBALL). Yezhou Wang was funded by the Fonds de recherche du Québec-Nature and technologies (FRQNT). This project/research/publication has received funding from the European Union’s Horizon Europe Programme under the Specific Grant Agreement No. 101147319 (EBRAINS 2.0 Project; KA). Dr. Nicole Eichert was supported by a Sir Henry Wellcome Postdoctoral Fellowship from the Wellcome Trust [222799/Z/21/Z]. Dr. Jessica Royer was supported by a fellowship from the Canadian Institute of Health Research (CIHR). Dr. Boris Bernhardt furthermore acknowledges research support from the National Science and Engineering Research Council of Canada (NSERC Discovery-1304413), the Canadian Institutes of Health Research (CIHR FDN-154298, PJT-174995, PJT-191853), SickKids Foundation (NI17-039), Azrieli Center for Autism Research (ACAR-TACC), BrainCanada (Future-Leaders), and the Tier-2 Canada Research Chairs (CRC) program.

## References

1. H.-J. Park, K. Friston, Structural and Functional Brain Networks: From Connections to Cognition. Science 342, 1238411 (2013).

2. D. J. Felleman, D. C. Van Essen, Distributed Hierarchical Processing in the Primate Cerebral Cortex. Cereb. Cortex 1, 1–47 (1991).

3. M. F. Glasser et al., A multi-modal parcellation of human cerebral cortex. Nature 536, 171–178 (2016).

4. D. S. Bassett, E. Bullmore, Small-World Brain Networks. Neuroscientist 12, 512–523 (2006).

5. E. Bullmore, O. Sporns, The economy of brain network organization. Nature Reviews Neuroscience 13, 336–349 (2012).

6. M. M. Mesulam, Large-scale neurocognitive networks and distributed processing for attention, language, and memory. Ann. Neurol. 28, 597–613 (1990).

7. M. M. Mesulam, From sensation to cognition. Brain 121, 1013–1052 (1998).

8. J. Sepulcre et al., The Organization of Local and Distant Functional Connectivity in the Human Brain. PLoS Comput. Biol. 6, e1000808 (2010).

9. G. L. Poerio et al., The role of the default mode network in component processes underlying the wandering mind. Soc. Cogn. Affect. Neurosci. 12, 1047–1062 (2017).

10. D. L. Schacter, R. G. Benoit, K. K. Szpunar, Episodic Future Thinking: Mechanisms and Functions. Curr Opin Behav Sci 17, 41–50 (2017).

11. M. Amft et al., Definition and characterization of an extended social-affective default network. Brain Structure and Function 220, 1031–1049 (2015).

12. R. L. Buckner, F. M. Krienen, The evolution of distributed association networks in the human brain. Trends Cogn Sci 17, 648–665 (2013).

13. F. M. Krienen, R. L. Buckner, “Chapter 35 - Human Association Cortex: Expanded, Untethered, Neotenous, and Plastic” in Evolutionary Neuroscience (Second Edition), J. H. Kaas, Ed. (Academic Press, London, 2020), 10.1016/B978-0-12-820584-6.00035-0, pp. 845–860.

14. K. Brodmann, Vergleichende Lokalisationslehre der Grosshirnrinde in ihren Prinzipien dargestellt auf Grund des Zellenbaues von Dr. K. Brodmann (J.A. Barth, 1909).

15. A. Schaefer et al., Local-Global Parcellation of the Human Cerebral Cortex from Intrinsic Functional Connectivity MRI. Cereb. Cortex 28, 3095–3114 (2018).

16. K. Amunts, H. Mohlberg, S. Bludau, K. Zilles, Julich-Brain: A 3D probabilistic atlas of the human brain’s cytoarchitecture. Science 369, 988 (2020).

17. K. Amunts, K. Zilles, Architectonic Mapping of the Human Brain beyond Brodmann. Neuron 88, 1086–1107 (2015).

18. F. Sanides, Grenzerscheinungen an myeloarchitektonischen Feldergrenzen. Dtsch. Z. Nervenheilkd. 180, 381–405 (1960).

19. D. S. Margulies et al., Situating the default-mode network along a principal gradient of macroscale cortical organization. Proc. Natl. Acad. Sci. U. S. A. 113, 12574–12579 (2016).

20. C. Paquola et al., Microstructural and functional gradients are increasingly dissociated in transmodal cortices. PLoS Biol. 17, e3000284 (2019).

21. C. Paquola et al., Shifts in myeloarchitecture characterise adolescent development of cortical gradients. Elife 8 (2019).

22. C. Paquola et al., Convergence of cortical types and functional motifs in the human mesiotemporal lobe. eLife 9, e60673 (2020).

23. J. M. Huntenburg et al., A Systematic Relationship Between Functional Connectivity and Intracortical Myelin in the Human Cerebral Cortex. Cereb. Cortex 27, 981–997 (2017).

24. S. L. Valk et al., Genetic and phylogenetic uncoupling of structure and function in human transmodal cortex. bioRxiv 10.1101/2021.06.08.447522, 2021.2006.2008.447522 (2021).

25. J. M. Huntenburg, P.-L. Bazin, D. S. Margulies, Large-Scale Gradients in Human Cortical Organization. Trends Cogn Sci 22, 21–31 (2018).

26. K. V. Haak, C. F. Beckmann, Understanding brain organisation in the face of functional heterogeneity and functional multiplicity. Neuroimage 220, 117061 (2020).

27. D. C. Van Essen, D. L. Dierker, Surface-Based and Probabilistic Atlases of Primate Cerebral Cortex. Neuron 56, 209–225 (2007).

28. J. Hill et al., Similar patterns of cortical expansion during human development and evolution. Proceedings of the National Academy of Sciences 107, 13135–13140 (2010).

29. B. T. Thomas Yeo et al., The organization of the human cerebral cortex estimated by intrinsic functional connectivity. J. Neurophysiol. 106, 1125–1165 (2011).

30. S. L. Valk et al., Genetic and phylogenetic uncoupling of structure and function in human transmodal cortex. Nat Commun 13, 2341 (2022).

31. S. L. Valk et al., Shaping brain structure: Genetic and phylogenetic axes of macroscale organization of cortical thickness. Science Advances 6, eabb3417.

32. R. M. Braga, R. Leech, Echoes of the Brain: Local-Scale Representation of Whole-Brain Functional Networks within Transmodal Cortex. Neuroscientist 21, 540–551 (2015).

33. J. Smallwood et al., Escaping the here and now: Evidence for a role of the default mode network in perceptually decoupled thought. Neuroimage 69, 120–125 (2013).

34. D. S. Margulies, J. Smallwood, Converging evidence for the role of transmodal cortex in cognition. Proceedings of the National Academy of Sciences 114, 12641–12643 (2017).

35. M. Konishi, D. G. McLaren, H. Engen, J. Smallwood, Shaped by the Past: The Default Mode Network Supports Cognition that Is Independent of Immediate Perceptual Input. PLoS One 10, e0132209 (2015).

36. S. Larivière et al., Microstructure-Informed Connectomics: Enriching Large-Scale Descriptions of Healthy and Diseased Brains. Brain Connectivity 9, 113–127 (2018).

37. R. F. Betzel, D. S. Bassett, Multi-scale brain networks. Neuroimage 160, 73–83 (2017).

38. N. A. Bock et al., Optimizing T1-weighted imaging of cortical myelin content at 3.0T. Neuroimage 65, 1–12 (2013).

39. J. Dinse et al. (2013) A Histology-Based Model of Quantitative T1 Contrast for In-vivo Cortical Parcellation of High-Resolution 7 Tesla Brain MR Images. in Medical Image Computing and Computer-Assisted Intervention – MICCAI 2013, eds K. Mori, I. Sakuma, Y. Sato, C. Barillot, N. Navab (Springer Berlin Heidelberg, Berlin, Heidelberg), pp 51–58.

40. C. Stüber et al., Myelin and iron concentration in the human brain: A quantitative study of MRI contrast. Neuroimage 93, 95–106 (2014).

41. C. Paquola et al., A multi-scale cortical wiring space links cellular architecture and functional dynamics in the human brain. PLoS Biol. 18, e3000979 (2020).

42. E. Bullmore, O. Sporns, Complex brain networks: graph theoretical analysis of structural and functional systems. Nat. Rev. Neurosci. 10, 186–198 (2009).

43. G. Gong et al., Mapping anatomical connectivity patterns of human cerebral cortex using in vivo diffusion tensor imaging tractography. Cerebral cortex (New York, N.Y.: 1991) 19, 524–536 (2009).

44. M. Mesulam, The evolving landscape of human cortical connectivity: Facts and inferences. Neuroimage 62, 2182–2189 (2012).

45. A. Goulas, K. Zilles, C. C. Hilgetag, Cortical Gradients and Laminar Projections in Mammals. Trends Neurosci. 41, 775–788 (2018).

46. E. Koechlin, C. Ody, F. Kouneiher, The Architecture of Cognitive Control in the Human Prefrontal Cortex. Science 302, 1181–1185 (2003).

47. D. Badre, M. D’Esposito, Is the rostro-caudal axis of the frontal lobe hierarchical? Nature Reviews Neuroscience 10, 659–669 (2009).

48. M. Visser, E. Jefferies, K. V. Embleton, M. A. Lambon Ralph, Both the Middle Temporal Gyrus and the Ventral Anterior Temporal Area Are Crucial for Multimodal Semantic Processing: Distortion-corrected fMRI Evidence for a Double Gradient of Information Convergence in the Temporal Lobes. J. Cogn. Neurosci. 24, 1766–1778 (2012).

49. D. S. Margulies et al., Mapping the functional connectivity of anterior cingulate cortex. Neuroimage 37, 579–588 (2007).

50. C. Kelly et al., A convergent functional architecture of the insula emerges across imaging modalities. Neuroimage 61, 1129–1142 (2012).

51. R. Vos de Wael et al., Anatomical and microstructural determinants of hippocampal subfield functional connectome embedding. Proceedings of the National Academy of Sciences 115, 10154 (2018).

52. S. M. Smith et al., Correspondence of the brain’s functional architecture during activation and rest. Proceedings of the National Academy of Sciences 106, 13040 (2009).

53. A. G. van der Kolk, J. Hendrikse, J. J. M. Zwanenburg, F. Visser, P. R. Luijten, Clinical applications of 7T MRI in the brain. Eur. J. Radiol. 82, 708–718 (2013).

54. C. Triantafyllou et al., Comparison of physiological noise at 1.5 T, 3 T and 7 T and optimization of fMRI acquisition parameters. Neuroimage 26, 243–250 (2005).

55. O. Kraff, A. Fischer, A. M. Nagel, C. Mönninghoff, M. E. Ladd, MRI at 7 tesla and above: Demonstrated and potential capabilities. J. Magn. Reson. Imaging 41, 13–33 (2015).

56. R. Sladky et al., High-resolution functional MRI of the human amygdala at 7T. Eur. J. Radiol. 82, 728–733 (2013).

57. W. van der Zwaag et al., fMRI at 1.5, 3 and 7 T: Characterising BOLD signal changes. Neuroimage 47, 1425–1434 (2009).

58. D. Schluppeck, R.-M. Sanchez-Panchuelo, S. T. Francis, Exploring structure and function of sensory cortex with 7T MRI. Neuroimage 164, 10–17 (2018).

59. K. Koller et al., MICRA: Microstructural image compilation with repeated acquisitions. Neuroimage 225, 117406 (2021).

60. Timothy O. Laumann et al., Functional System and Areal Organization of a Highly Sampled Individual Human Brain. Neuron 87, 657–670 (2015).

61. L. E. Maxwell et al., Precision Brain Morphometry Using Cluster Scanning. medRxiv 10.1101/2023.12.23.23300492, 2023.2012.2023.23300492 (2023).

62. J. W. Cho, A. Korchmaros, J. T. Vogelstein, M. P. Milham, T. Xu, Impact of concatenating fMRI data on reliability for functional connectomics. Neuroimage 226, 117549 (2021).

63. E. M. Gordon et al., Precision Functional Mapping of Individual Human Brains. Neuron 95, 791-807.e797 (2017).

64. J. S. Elam et al., The Human Connectome Project: A retrospective. Neuroimage 244, 118543 (2021).

65. B.-y. Park et al., Differences in subcortico-cortical interactions identified from connectome and microcircuit models in autism. Nat Commun 12, 2225 (2021).

66. Y. Wang et al., Long-range functional connections mirror and link microarchitectural and cognitive hierarchies in the human brain. Cereb. Cortex 33, 1782–1798 (2023).

67. J. Royer et al., An Open MRI Dataset For Multiscale Neuroscience. Scientific Data 9, 569 (2022).

68. R. Vos de Wael et al., BrainSpace: a toolbox for the analysis of macroscale gradients in neuroimaging and connectomics datasets. Commun Biol 3, 103 (2020).

69. A. F. Alexander-Bloch et al., On testing for spatial correspondence between maps of human brain structure and function. Neuroimage 178, 540–551 (2018).

70. M. M. Mesulam, Principles of Behavioral and Cognitive Neurology (Oxford University Press, 2000).

71. K. Amunts et al., BigBrain: An Ultrahigh-Resolution 3D Human Brain Model. Science 340, 1472 (2013).

72. P. Taylor, J. N. Hobbs, J. Burroni, H. T. Siegelmann, The global landscape of cognition: hierarchical aggregation as an organizational principle of human cortical networks and functions. Sci. Rep. 5, 18112 (2015).

73. S. Zeki, S. Shipp, The functional logic of cortical connections. Nature 335, 311–317 (1988).

74. T. E. J. Behrens, O. Sporns, Human connectomics. Curr. Opin. Neurobiol. 22, 144–153 (2012).

75. J. Bijsterbosch et al., Challenges and future directions for representations of functional brain organization. Nat. Neurosci. 23, 1484–1495 (2020).

76. K. J. Friston, Modalities, Modes, and Models in Functional Neuroimaging. Science 326, 399–403 (2009).

77. C. Paquola, K. Amunts, A. Evans, J. Smallwood, B. Bernhardt, Closing the mechanistic gap: the value of microarchitecture in understanding cognitive networks. Trends Cogn Sci 26, 873–886 (2022).

78. P. Balchandani, T. P. Naidich, Ultra-High-Field MR Neuroimaging. Am. J. Neuroradiol. 36, 1204 (2015).

79. H.-M. Dong, D. S. Margulies, X.-N. Zuo, A. J. Holmes, Shifting gradients of macroscale cortical organization mark the transition from childhood to adolescence. Proceedings of the National Academy of Sciences 118, e2024448118 (2021).

80. J. B. Burt et al., Hierarchy of transcriptomic specialization across human cortex captured by structural neuroimaging topography. Nat. Neurosci. 21, 1251–1259 (2018).

81. E. T. Rolls, Limbic systems for emotion and for memory, but no single limbic system. Cortex 62, 119–157 (2015).

82. P. E. Vértes et al., Gene transcription profiles associated with inter-modular hubs and connection distance in human functional magnetic resonance imaging networks. Philosophical Transactions of the Royal Society B: Biological Sciences 371, 20150362 (2016).

83. J. D. Schmahmann, D. N. Pandya (2006) Fiber Pathways of the Brain. (Oxford University Press).

84. J. Richiardi et al., Correlated gene expression supports synchronous activity in brain networks. Science 348, 1241 (2015).

85. H. Barbas, Pattern in the laminar origin of corticocortical connections. J. Comp. Neurol. 252, 415–422 (1986).

86. H. Barbas, N. Rempel-Clower, Cortical structure predicts the pattern of corticocortical connections. Cereb. Cortex 7, 635–646 (1997).

87. H. Barbas, General Cortical and Special Prefrontal Connections: Principles from Structure to Function. Annu. Rev. Neurosci. 38, 269–289 (2015).

88. G. Rothschild, A. Mizrahi, Global Order and Local Disorder in Brain Maps. Annu. Rev. Neurosci. 38, 247–268 (2015).

89. S.-C. Yen, J. Baker, C. M. Gray, Heterogeneity in the Responses of Adjacent Neurons to Natural Stimuli in Cat Striate Cortex. J. Neurophysiol. 97, 1326–1341 (2007).

90. C. D. Gregory, M. G. Geoffrey, O. Izumi, D. F. Ralph, Functional Micro-Organization of Primary Visual Cortex: Receptive Field Analysis of Nearby Neurons. J Neurosci 19, 4046 (1999).

91. D. S. Reich, F. Mechler, J. D. Victor, Independent and Redundant Information in Nearby Cortical Neurons. Science 294, 2566–2568 (2001).

92. A. C. M. Kevan, S. Sylvia, Functional Heterogeneity in Neighboring Neurons of Cat Primary Visual Cortex in Response to Both Artificial and Natural Stimuli. J Neurosci 33, 7325 (2013).

93. G. Chechik et al., Reduction of Information Redundancy in the Ascending Auditory Pathway. Neuron 51, 359–368 (2006).

94. P. Casey et al., The Unique Cytoarchitecture and Wiring of the Human Default Mode Network. bioRxiv 10.1101/2021.11.22.469533, 2021.2011.2022.469533 (2021).

95. J. H. Kaas, Evolution of the neocortex. Curr. Biol. 16, R910–R914 (2006).

96. L. Audrey et al., Functional Connectivity Development along the Sensorimotor-Association Axis Enhances the Cortical Hierarchy. bioRxiv 10.1101/2023.07.20.549090, 2023.2007.2020.549090 (2023).

97. V. Rajmohan, E. Mohandas, The limbic system. Indian journal of psychiatry 49, 132–139 (2007).

98. L. Chanes, L. F. Barrett, Redefining the Role of Limbic Areas in Cortical Processing. Trends Cogn Sci 20, 96–106 (2016).

99. R. F. Betzel et al., The community structure of functional brain networks exhibits scale-specific patterns of inter- and intra-subject variability. Neuroimage 202, 115990 (2019).

100. R. A. M. Haast, D. Ivanov, E. Formisano, K. Uludag, Reproducibility and Reliability of Quantitative and Weighted T(1) and T(2)(*) Mapping for Myelin-Based Cortical Parcellation at 7 Tesla. Front. Neuroanat. 10, 112–112 (2016).

101. J. P. Marques et al., MP2RAGE, a self bias-field corrected sequence for improved segmentation and T1-mapping at high field. Neuroimage 49, 1271–1281 (2010).

102. X. Li, P. S. Morgan, J. Ashburner, J. Smith, C. Rorden, The first step for neuroimaging data analysis: DICOM to NIfTI conversion. J. Neurosci. Methods 264, 47–56 (2016).

103. K. J. Gorgolewski et al., The brain imaging data structure, a format for organizing and describing outputs of neuroimaging experiments. Scientific data 3, 160044–160044 (2016).

104. R. R. Cruces et al., Micapipe: A pipeline for multimodal neuroimaging and connectome analysis. Neuroimage 263, 119612 (2022).

105. M. Jenkinson, C. F. Beckmann, T. E. J. Behrens, M. W. Woolrich, S. M. Smith, FSL. Neuroimage 62, 782–790 (2012).

106. B. B. Avants et al., The optimal template effect in hippocampus studies of diseased populations. Neuroimage 49, 2457–2466 (2010).

107. S. M. Smith, Fast robust automated brain extraction. Hum. Brain Mapp. 17, 143–155 (2002).

108. L. Henschel et al., FastSurfer - A fast and accurate deep learning based neuroimaging pipeline. Neuroimage 219, 117012 (2020).

109. J. D. Tournier et al., MRtrix3: A fast, flexible and open software framework for medical image processing and visualisation. Neuroimage 202, 116137 (2019).

110. J. Veraart et al., Denoising of diffusion MRI using random matrix theory. Neuroimage 142, 394–406 (2016).

111. L. Cordero-Grande, D. Christiaens, J. Hutter, A. N. Price, J. V. Hajnal, Complex diffusion-weighted image estimation via matrix recovery under general noise models. Neuroimage 200, 391–404 (2019).

112. R. E. Smith, J.-D. Tournier, F. Calamante, A. Connelly, Anatomically-constrained tractography: Improved diffusion MRI streamlines tractography through effective use of anatomical information. Neuroimage 62, 1924–1938 (2012).

113. B. B. Avants, C. L. Epstein, M. Grossman, J. C. Gee, Symmetric diffeomorphic image registration with cross-correlation: Evaluating automated labeling of elderly and neurodegenerative brain. Med. Image Anal. 12, 26–41 (2008).

114. R. W. Cox, AFNI: Software for Analysis and Visualization of Functional Magnetic Resonance Neuroimages. Comput. Biomed. Res. 29, 162–173 (1996).

115. P. Kundu et al., Integrated strategy for improving functional connectivity mapping using multiecho fMRI. Proceedings of the National Academy of Sciences 110, 16187–16192 (2013).

116. P. Kundu, S. J. Inati, J. W. Evans, W.-M. Luh, P. A. Bandettini, Differentiating BOLD and non-BOLD signals in fMRI time series using multi-echo EPI. Neuroimage 60, 1759–1770 (2012).

117. L. Lemieux, A. Salek-Haddadi, T. E. Lund, H. Laufs, D. Carmichael, Modelling large motion events in fMRI studies of patients with epilepsy. Magn. Reson. Imaging 25, 894–901 (2007).

118. T. D. Satterthwaite et al., An improved framework for confound regression and filtering for control of motion artifact in the preprocessing of resting-state functional connectivity data. Neuroimage 64, 240–256 (2013).

119. D. N. Greve, B. Fischl, Accurate and robust brain image alignment using boundary-based registration. Neuroimage 48, 63–72 (2009).

120. D. Marcus et al., Informatics and Data Mining Tools and Strategies for the Human Connectome Project. 5 (2011).

121. D. C. Van Essen, M. F. Glasser, D. L. Dierker, J. Harwell, T. Coalson, Parcellations and Hemispheric Asymmetries of Human Cerebral Cortex Analyzed on Surface-Based Atlases. Cereb. Cortex 22, 2241–2262 (2012).

122. J. Royer et al., Myeloarchitecture gradients in the human insula: Histological underpinnings and association to intrinsic functional connectivity. Neuroimage 216, 116859 (2020).

123. M. D. Waehnert et al., Anatomically motivated modeling of cortical laminae. Neuroimage 93, 210–220 (2014).

124. D. Christiaens et al., Global tractography of multi-shell diffusion-weighted imaging data using a multi-tissue model. Neuroimage 123, 89–101 (2015).

125. B. Jeurissen, J.-D. Tournier, T. Dhollander, A. Connelly, J. Sijbers, Multi-tissue constrained spherical deconvolution for improved analysis of multi-shell diffusion MRI data. Neuroimage 103, 411–426 (2014).

126. J. D. Tournier, F. Calamante, D. G. Gadian, A. Connelly, Direct estimation of the fiber orientation density function from diffusion-weighted MRI data using spherical deconvolution. Neuroimage 23, 1176–1185 (2004).

127. R. E. Smith, J.-D. Tournier, F. Calamante, A. Connelly, SIFT2: Enabling dense quantitative assessment of brain white matter connectivity using streamlines tractography. Neuroimage 119, 338–351 (2015).

128. R. R. Coifman et al., Geometric diffusions as a tool for harmonic analysis and structure definition of data: multiscale methods. Proc. Natl. Acad. Sci. U. S. A. 102, 7432–7437 (2005).

