## SUPPLEMENTARY MATERIALS for "MULTIMODAL GRADIENTS UNIFY LOCAL AND GLOBAL CORTICAL ORGANIZATION"

#### SUPPLEMENTARY METHODS

##### *MRI acquisition*

*MICA-MICs.* Two T1w scans with identical parameters were acquired with a 3D magnetization-prepared rapid gradient echo (MPRAGE) sequence (0.8mm isovoxels, matrix=320×320, 224 sagittal slices, repetition time (TR)=2300 ms, echo time (TE)=3.14 ms, inversion time (TI)=900 ms, flip angle=9°, iPAT=2). Scans were visually examined to ensure minimal head motion, and repeated if necessary.

qT1 relaxometry data was acquired using a 3D-MP2RAGE sequence (0.8mm isotropic, 240 sagittal slices, TR=5000 ms, TE=2.9ms, TI 1=940 ms, T1 2=2830 ms, flip angle=4°, flip angle 2=5°, iPAT=3, bandwidth=270 Hz/px, echo spacing=7.2ms, partial Fourier=6/8). Both inversion images were combined for qT1 mapping to minimize sensitivity to B1 inhomogeneities and optimize intra- and inter-subject reliability [1,2].

DWI data was acquired using a 2D spin-echo echo-planar imaging sequence, consisting of three shells with b-values=300, 700, and 2000s/mm$^2$, and with 10, 40, and 90 diffusion weighting directions per shell, respectively (1.6 mm isotropic voxels, TR = 3500 ms, TE = 64.40 ms, flip angle = 90°, refocusing flip angle = 180°, FOV = 224 × 224 mm$^2$, slice thickness = 1.6 mm, multi-band factor = 3, echo spacing = 0.76 ms). b0 images acquired in reverse phase encoding direction were used for distortion correction of DWI scans.

A 7-minute rs-fMRI scan was acquired using multiband accelerated 2D-BOLD echo-planar imaging (3 mm isovoxels, TR=600 ms, TE=30 ms, flip angle=52°, FOV=240×240 mm$^2$, slice thickness=3 mm, mb factor=6, echo spacing=0.54 ms). Participants were instructed to keep their eyes open, not fall asleep, and look at a fixation cross. Two spin-echo images with reverse phase encoding were also included for distortion correction of the rs-fMRI scans (phase encoding=AP/PA, 3mm isovoxels, TR=4029 ms, TE=48 ms, flip angle=90°, FOV=240×240 mm$^2$, slice thickness=3mm, echo spacing=0.54 ms, bandwidth= 2084 Hz/Px).

##### *Multimodal MRI processing*

*MICA-MICs.* The surface reconstructions for each subject underwent manual inspection and correction for segmentation errors by placing control points and applying manual edits. The qT1 scans were linearly co-registered to the corresponding subject's T1w scan.

Regarding the DWI data, pre-processing was carried out using MRtrix [3] in the native DWI space. The DWI data underwent denoising [4,5], b0 intensity normalization, and correction for susceptibility distortion, head motion, and eddy currents. These corrections were performed using FSL and involved utilizing two b=0 s/mm$^2$ volumes with reverse phase encoding. Anatomical masks for tractography were non-linearly co-registered to native DWI space using the deformable SyN approach implemented in ANTs [6].

For the rs-fMRI scans, pre-processing steps were conducted using AFNI [7] and FSL[8] tools. The first five volumes were discarded to ensure magnetic field saturation. The remaining volumes underwent reorientation, motion correction, and distortion correction. We applied FMRIB's ICA-based X-noiseifier (ICA-FIX) [9] and spike regression to remove timepoints with large motion spikes, effectively removing nuisance signals [10,11]. The volume timeseries were registered to FastSurfer [12] space using boundary-based registration implemented in ANTs using linear and non-linear methods [13].

#### SUPPLEMENTARY RESULTS

*Homogeneity across parcels*

Differences between cortical regions are crucial for their functional specialization, while their similarities support functional integration. To examine cortical homogeneity, we calculated '1 - cosine distance' to represent the similarity between regional gradient profiles, resulting in an affinity matrix (**Figure S1A**). We estimated the parcel-wise similarity coefficient by taking the mean value of each row in the affinity matrix and examined its distribution. Note that this computation is not simply the inverse of the heterogeneity. The analysis revealed that the idiotypic system exhibited the lowest similarity, while the paralimbic system showed the highest similarity. Hierarchical clustering on the affinity matrix resulted in a similar result (**Figure S1B**), providing four robust clusters that recapitulated sensory-fugal hierarchies (**Figure S1C).**

*Patterns of intra-parcel assessments.*

In addition to examining relationships between cortical parcels, exploring the layout within each parcel helps us to better understand local cortical organization. To validate our hypothesis that vertices within a parcel are more homogeneous, but different patterns may exist in specific parcels, we investigated intra-parcel heterogeneity at the vertex level. For each vertex, we defined its local dissimilarity as cosine distance between its gradients and the gradient profiles of the parcel to which it belongs (**Figure S2A**). Within a parcel, this measure characterizes the difference between the gradients of individual vertices and the average of all gradients. Local dissimilarity was found considerably lower compared to the global dissimilarity, providing support for our hypothesis. Interestingly, we identified regions with higher local dissimilarity, such as pre-SMA and SPL. We found that the heteromodal and unimodal systems exhibited higher local dissimilarity ($p_{spin}<0.05$, FDR correction, **Figure S2B**), indicating a distinct pattern of multimodal gradients within parcels of these systems. In contrast, paralimbic systems showed lower local dissimilarity ($p_{spin}<0.05$, FDR correction). Upon examining associations between histological gradient and local dissimilarity, we observed a trend toward a negative correlation (rho=-0.17, $p_{spin}=0.12$).

### FIGURES

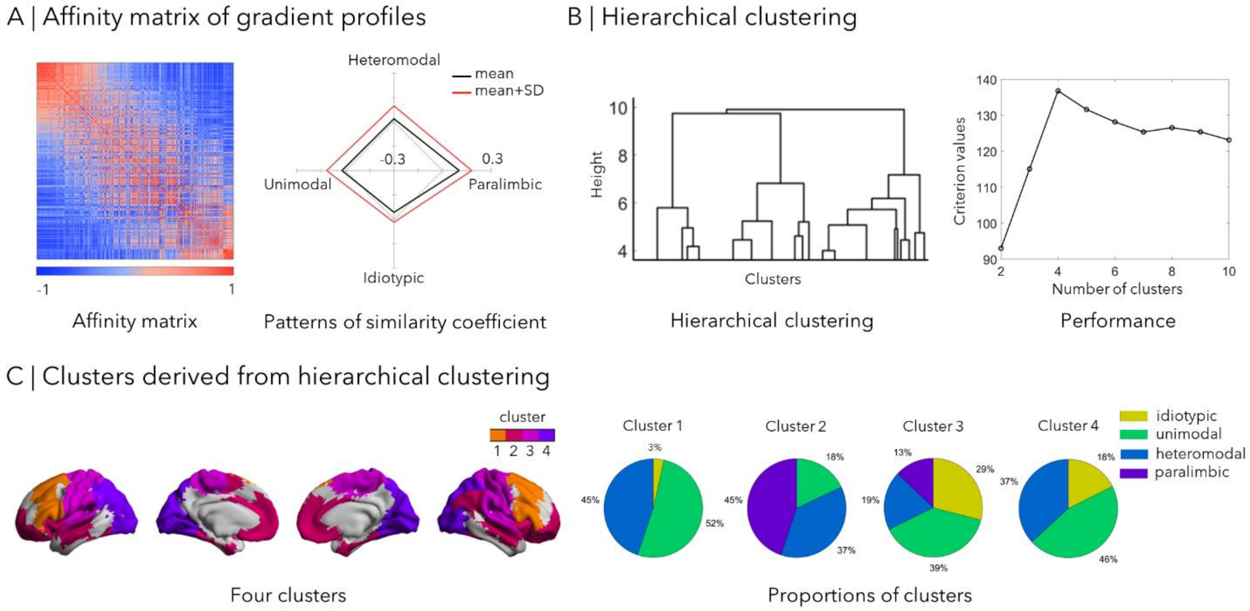


**Figure S1. Homogenous patterns of cortical parcels. (A) Affinity matrix of gradient profiles of MPC, SC and FC**. We constructed an affinity matrix by calculating the cosine similarity between cortical parcels. The similarity coefficient was determined by averaging each row of the affinity matrix. We examined this coefficient in four cortical hierarchies. **(B) Hierarchical clustering**. We performed hierarchical clustering based on the affinity matrix. The left panel illustrates the clustering results, while the right panel displays the criterion values of hierarchical clustering with different cluster numbers. **(C) Clusters derived from hierarchical clustering**. The left panel showcases the four clusters obtained from hierarchical clustering. The right panel shows the proportions of four cortical hierarchies within each cluster.

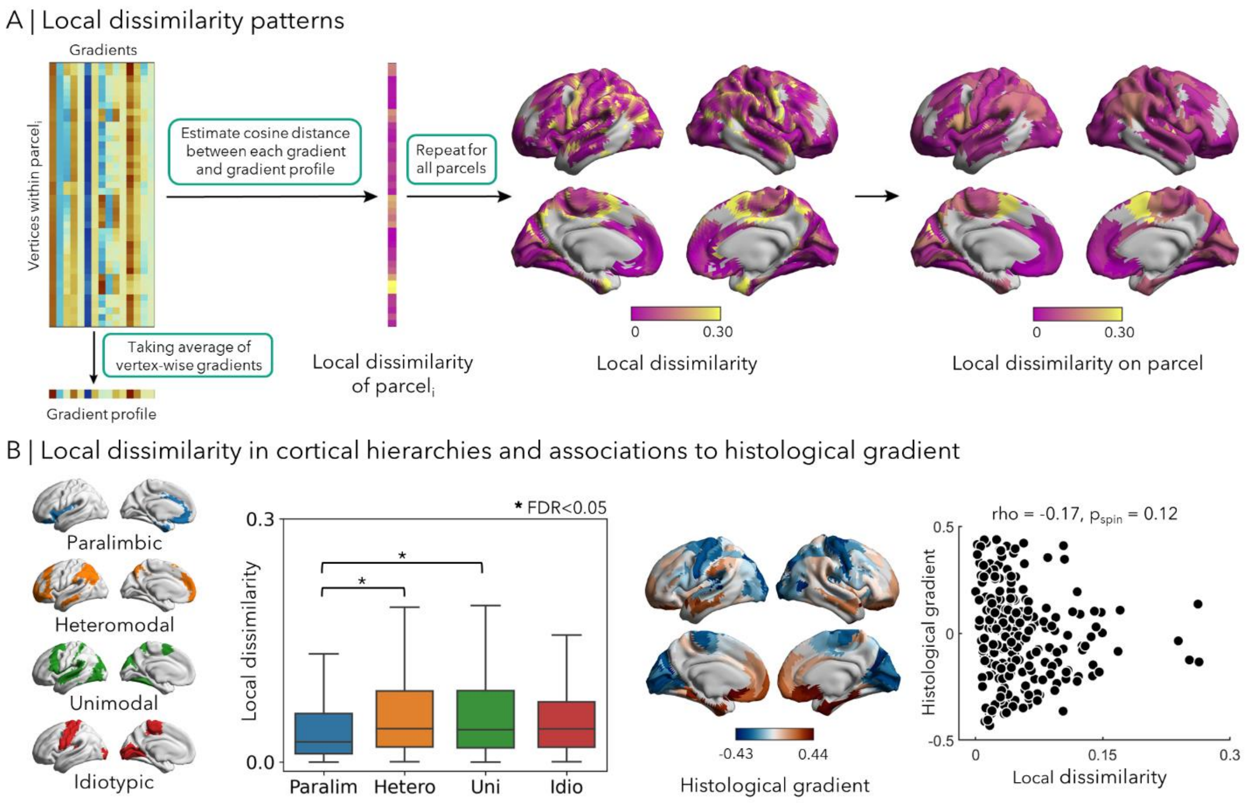


**Figure S2. Intra-parcel heterogeneity. (A) Local dissimilarity patterns**. The gradient profile of parcel *i* was calculated by averaging the gradients across all vertices belonging to parcel *i*. The local dissimilarity of parcel *i* was then determined by calculating cosine distance between vertex-wise gradients and gradient profile of parcel *i*. By repeating this procedure for all parcels, we generated a map illustrating the distribution of local dissimilarity across the cortex. **(B) Local dissimilarity in cortical hierarchies and associations to histological gradient**. The distribution of local dissimilarity in four cortical hierarchies is shown in the left panel. To assess differences between each cortical hierarchy, we performed two-sample t-tests with FDR correction. To explore associations with the histological gradient, Spearman's correlation coefficient was computed with FDR correction.

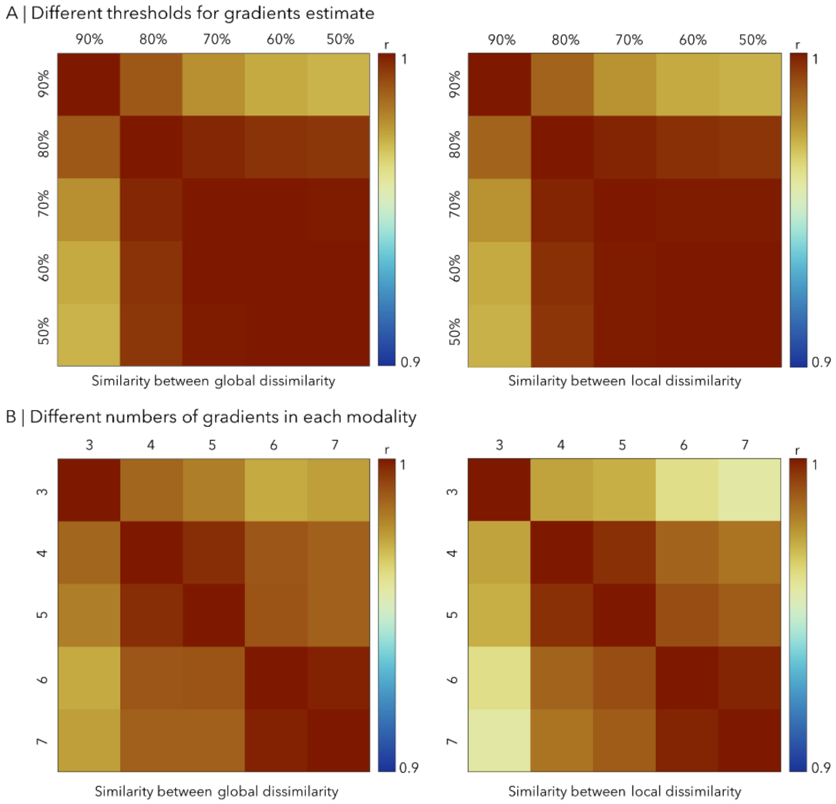


**Figure S3. Robustness analysis. (A) Different thresholds for gradients estimate.** Multimodal gradients were recalculated using connectivity thresholds ranging from 50% to 90%. Pearson's correlation coefficients were computed between global/local dissimilarities derived from multimodal gradients with varying thresholds. **(B) Different numbers of gradients in each modality.** Gradient profiles were recalculated using different numbers of gradients (3-7). Pearson's correlation coefficients were computed between global/local dissimilarity derived from gradient profiles estimated with varying numbers of gradients.

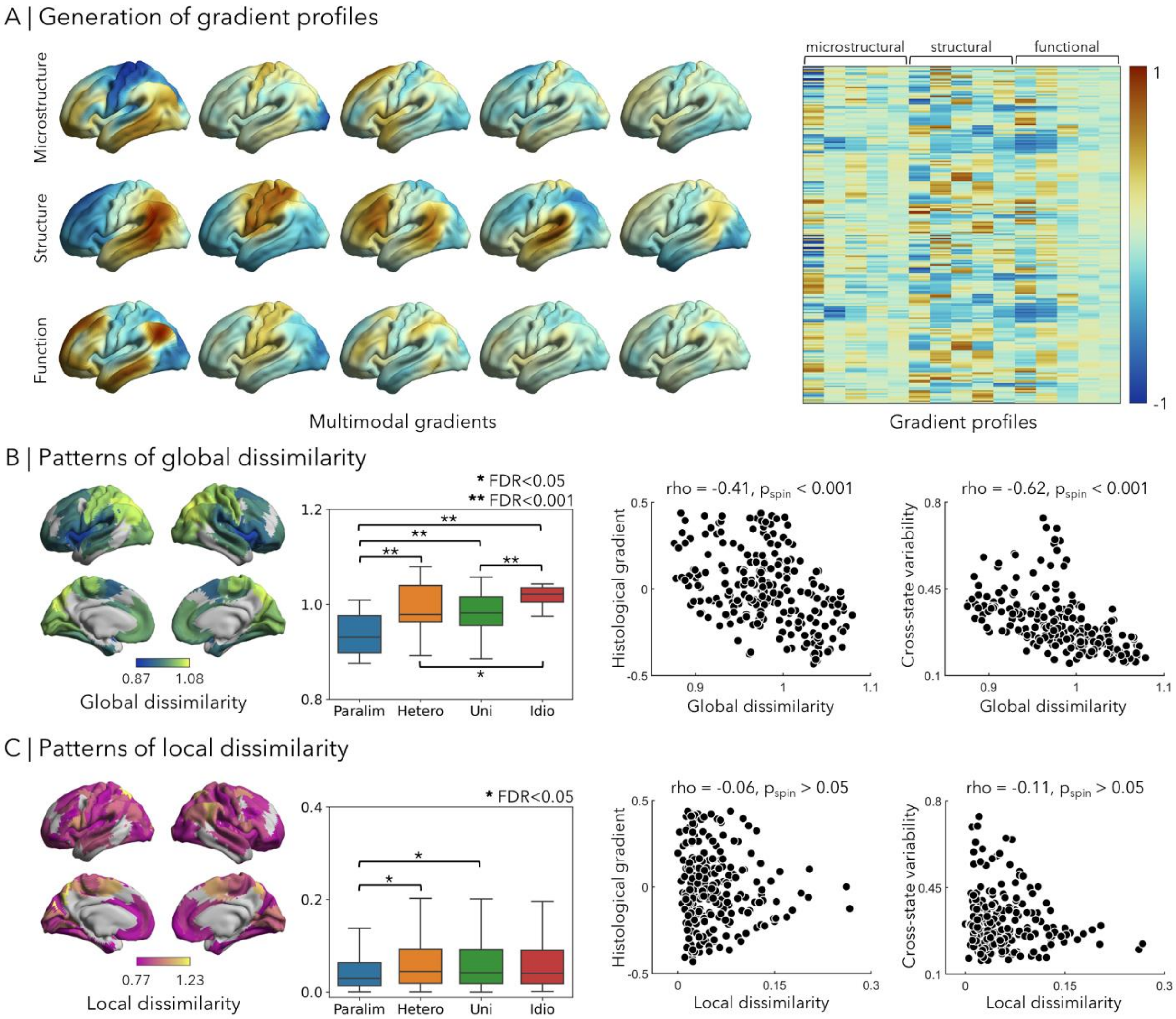


**Figure S4. Replication analysis based on 3T MRI. (A) Generation of gradient profiles**. Multimodal gradients and gradient profiles were generated based on 3T MRI data. **(B) Patterns of global dissimilarity.** Global dissimilarity was estimated based on 3T MRI data. To discern distinctions between each cortical hierarchy, two-sample t-tests were conducted with FDR correction. Spearman's correlation coefficients were computed to examine associations between global dissimilarity and histological gradient, as well as cross-state variability, with p-values corrected using spin permutation tests. **(C) Patterns of local dissimilarity.** Local dissimilarity was estimated based on 3T MRI data. To discern distinctions between each cortical hierarchy, two-sample t-tests were conducted with FDR correction. Spearman's correlation coefficients were computed to examine associations between local dissimilarity and histological gradient, as well as cross-state variability.

#### Supplemental References


1 Haast, R. A. M., Ivanov, D., Formisano, E. & Uludağ, K. Reproducibility and Reliability of Quantitative and Weighted T(1) and T(2)(∗) Mapping for Myelin-Based Cortical Parcellation at 7 Tesla. *Front. Neuroanat.* **10**, 112-112, doi:10.3389/fnana.2016.00112 (2016).

2 Marques, J. P. *et al.* MP2RAGE, a self bias-field corrected sequence for improved segmentation and T1-mapping at high field. *Neuroimage* **49**, 1271-1281, doi:https://doi.org/10.1016/j.neuroimage.2009.10.002 (2010).

3 Tournier, J. D. *et al.* MRtrix3: A fast, flexible and open software framework for medical image processing and visualisation. *Neuroimage* **202**, 116137, doi:https://doi.org/10.1016/j.neuroimage.2019.116137 (2019).

4 Veraart, J. *et al.* Denoising of diffusion MRI using random matrix theory. *Neuroimage* **142**, 394-406, doi:https://doi.org/10.1016/j.neuroimage.2016.08.016 (2016).

5 Cordero-Grande, L., Christiaens, D., Hutter, J., Price, A. N. & Hajnal, J. V. Complex diffusion-weighted image estimation via matrix recovery under general noise models. *Neuroimage* **200**, 391-404, doi:https://doi.org/10.1016/j.neuroimage.2019.06.039 (2019).

6 Avants, B. B., Epstein, C. L., Grossman, M. & Gee, J. C. Symmetric diffeomorphic image registration with cross-correlation: Evaluating automated labeling of elderly and neurodegenerative brain. *Med. Image Anal.* **12**, 26-41, doi:https://doi.org/10.1016/j.media.2007.06.004 (2008).

7 Cox, R. W. AFNI: Software for Analysis and Visualization of Functional Magnetic Resonance Neuroimages. *Comput. Biomed. Res.* **29**, 162-173, doi:https://doi.org/10.1006/cbmr.1996.0014 (1996).

8 Jenkinson, M., Beckmann, C. F., Behrens, T. E. J., Woolrich, M. W. & Smith, S. M. FSL. *Neuroimage* **62**, 782-790, doi:https://doi.org/10.1016/j.neuroimage.2011.09.015 (2012).

9 Salimi-Khorshidi, G. *et al.* Automatic denoising of functional MRI data: Combining independent component analysis and hierarchical fusion of classifiers. *Neuroimage* **90**, 449-468, doi:https://doi.org/10.1016/j.neuroimage.2013.11.046 (2014).

10 Lemieux, L., Salek-Haddadi, A., Lund, T. E., Laufs, H. & Carmichael, D. Modelling large motion events in fMRI studies of patients with epilepsy. *Magn. Reson. Imaging* **25**, 894-901, doi:https://doi.org/10.1016/j.mri.2007.03.009 (2007).

11 Satterthwaite, T. D. *et al.* An improved framework for confound regression and filtering for control of motion artifact in the preprocessing of resting-state functional connectivity data. *Neuroimage* **64**, 240-256, doi:https://doi.org/10.1016/j.neuroimage.2012.08.052 (2013).

12 Henschel, L. *et al.* FastSurfer - A fast and accurate deep learning based neuroimaging pipeline. *Neuroimage* **219**, 117012, doi:https://doi.org/10.1016/j.neuroimage.2020.117012 (2020).

13 Greve, D. N. & Fischl, B. Accurate and robust brain image alignment using boundary-based registration. *Neuroimage* **48**, 63-72, doi:https://doi.org/10.1016/j.neuroimage.2009.06.060 (2009).
